# Dynamic single-molecule binding of carbamazepine explains diffuse cryo-EM density in the SUR1 drug-binding pocket

**DOI:** 10.64898/2026.06.13.732016

**Authors:** Christina Karafyllia, Katarzyna Walczewska-Szewc

## Abstract

Carbamazepine (CBZ), an anticonvulsant drug and pharmacochaperone of pancreatic ATP-sensitive potassium (KATP) channels, binds within the SUR1 drug-binding pocket. However, cryo-electron microscopy (cryo-EM) studies revealed an unusually diffuse ligand-associated density that could not be unambiguously interpreted. Despite being substantially smaller than glibenclamide (GBM), CBZ produced a density of comparable size, leading to the proposal that either two CBZ molecules occupy the binding pocket simultaneously or that a single molecule adopts multiple binding configuresions.

Here, we combine molecular dynamics simulations, density reconstruction from simulation ensembles, quantum chemical calculations, and binding free-energy analyses to investigate the molecular origin of this diffuse cryo-EM density. Density functional theory calculations indicate that CBZ dimers are only weakly stabilized and do not exhibit a strong intrinsic preference for dimerization. Molecular dynamics simulations further show that two simultaneously bound CBZ molecules fail to form a stable dimeric complex and generate density distributions inconsistent with the experimental cryo-EM map. In contrast, a single CBZ molecule remains localized within the SUR1 cavity while sampling a broad ensemble of positions and orientations. Remarkably, density reconstructed directly from the simulation trajectories closely reproduces the extent and shape of the experimentally observed cryo-EM density.

Our results demonstrate that the diffuse cryo-EM density associated with CBZ arises from dynamic binding of a single molecule rather than ligand dimerization. More broadly, this study highlights how ligand dynamics can shape cryo-EM densities and illustrates the value of integrating molecular simulations with experimental structural data to interpret heterogeneous ligand-binding states.

## 1. Introduction

Pancreatic ATP-sensitive potassium (K_ATP_) channels play a crucial role in glucose homeostasis by coupling glucose metabolism to insulin release. This channel functions as an intricate metabolic sensor. Each K_ATP_ channel is a hetero-octamer composed of four inward-rectifier potassium channel subunits (Kir6.2) forming the central pore and four surrounding regulatory sulfonylurea receptor subunits (SUR1)^1–3^. While SUR1 may lack intrinsic transport activity, it is indispensable for the proper trafficking, expression, and allosteric regulation of Kir6.2^2^. Together, these subunits regulate K+ efflux in response to changing intracellular ATP and ADP concentrations, thus controlling the resting membrane potential of pancreatic β-cells.

Mutations in pancreatic K_ATP_ channels are implicated in several diseases, most notably congenital hyperinsulinism and neonatal diabetes^4,5^. Pharmacological modulation of these channels is a core strategy in diabetes therapy. Typical anti-diabetic drugs, such as sulfonylureas like glibenclamide (GBM) and glinides, as well as the anticonvulsant carbamazepine (CBZ), are known channel inhibitors^6,7^. Although chemically diverse, these ligands exhibit functional similarity: they potently inhibit K_ATP_ channel activity by reducing the channel-open probability and abolishing the stimulatory effect of MgADP^6,8^. They may also act as pharmacochaperones, enabling proper channel folding and trafficking to the cell membrane. Functional and structural studies consistently indicate that these ligands occupy a common, high-affinity binding pocket within the SUR1 subunit, primarily located at the interface between the SUR1 transmembrane domains^6^.

Recent structural data further highlight the involvement of the N-terminal tail of the adjacent Kir6.2 subunit (KNtp) in this ligand-binding pocket^6,9^. This 30-amino-acid stretch of Kir6.2 inserts into the SUR1 core cavity, located immediately adjacent to the drug-binding site. This insertion correlates with a stabilized channel-closed state and is thought to be a key component of the mechanism for channel inhibition. Crucially, cryo-electron microscopy (cryo-EM) experiments have demonstrated a greater density corresponding to KNtp in the presence of both ATP and inhibitory ligands^6^. This suggests that CBZ and other inhibitors may stabilize the Kir6.2 N-terminus within the SUR1 ABC core, thereby further impeding channel opening and activity.

Cryo-EM studies by Martin et al.^6^ successfully resolved several inhibitors within the SUR1 binding pocket, including GBM and others. In the case of CBZ, the observed cryo-EM density did not allow unambiguous determination of the ligand’s configuration in the binding site. Surprisingly, despite CBZ having a molecular weight roughly half that of GBM, its observed cryo-EM density in the binding site was strikingly similar in size and shape to that of GBM. In the absence of high-resolution comparative structures, the authors proposed two primary, mutually exclusive explanations for this unexpected observation: either CBZ binds as a dimer, thus occupying the entire pocket, or it exhibits multiple, distinct binding poses (conformations), whose density is averaged out in the cryo-EM map.

The dimerization hypothesis is chemically plausible because carbamazepine is known to form stable intermolecular assemblies in both solution and crystalline phases^10^. Previous studies have shown that CBZ can adopt several dimeric arrangements stabilized by aromatic stacking or hydrogen-bonding interactions, depending on the surrounding environment^10,11^. However, whether such dimers can remain stable within the confined environment of the SUR1 cavity remains unknown. Equally plausible is the alternative possibility that the diffuse cryo-EM density arises from a dynamically bound single molecule that samples multiple orientations and positions while remaining localized within the binding pocket.

Molecular dynamics (MD) simulations provide a natural framework for addressing this ambiguity because they capture the ensemble of molecular configurations sampled by a ligand in its binding environment. Furthermore, simulation trajectories can be converted into three-dimensional spatial probability densities, enabling direct comparison between computationally sampled ligand distributions and experimentally observed cryo-EM densities. Such an approach allows not only evaluation of candidate binding modes but also assessment of whether the apparent size and shape of an experimental density can emerge from ligand dynamics alone.

Here, we combine molecular dynamics simulations, density reconstruction from simulation ensembles, quantum chemical calculations, and binding free-energy analyses to investigate the molecular origin of the diffuse cryo-EM density associated with carbamazepine in the SUR1 drug-binding pocket. Specifically, we test whether the experimental observations are more consistent with simultaneous binding of a CBZ dimer or with a dynamically bound single CBZ molecule sampling multiple poses. Our results demonstrate that the experimentally observed density can be explained by the latter mechanism and reveal a highly dynamic binding mode that distinguishes carbamazepine from the more rigidly bound inhibitor glibenclamide.

## 2. Methodology

### 2.1. System preparation and molecular dynamics simulations

For the analysis of CBZ dynamics within pancreatic KATP channels, two distinct systems were prepared, containing 1 and 2 CBZ ligands respectively. In both cases, we utilized the PDB entry 7u24, which consists of a Kir6.2-SUR1 pair, with the KNtp terminus and one GBM ligand positioned inside the Kir cavity^12^. This was also the reference system for our simulations.

Our structure was initially pre-processed using the Protein Preparation Wizard in Schrödinger (Version 11.7.012), which involved filling in missing side chains and determining their protonation states at pH 7. The structure contained several intrinsically disordered regions (IDRs), specifically residues 621-678, 742-766, 928-986, and 1044-1060. These segments were excluded from the model to maintain computational efficiency; preliminary structural assessments indicated that their omission did not adversely impact the conformational stability or the local dynamics of the target binding domains.

Subsequently, one and two CBZ molecules (Pubchem^13^, 3D conformer) were placed in the SUR1 drug-binding pocket using molecular docking. The highest-ranked pose obtained for each system was used as the starting configuration for MD simulations, yielding docking scores of -6.188 and -5.529 kcal mol⁻¹ for the single- and double-CBZ systems, respectively. Since the objective of docking was solely to generate physically reasonable initial configurations, all analyses and conclusions reported herein are derived from the MD trajectories rather than from docking results.

The simulation systems were subsequently generated using CHARMM-GUI^14^. This involved embedding the protein-ligand complexes within a POPC lipid bilayer and solvating them with a water box containing 0.15 M KCl ions, resulting in simulation boxes with approximate dimensions of 165 Å x 165 Å x 160 Å.

Equilibration of both systems was performed in GROMACS 2023^15^ through a stepwise release of positional restraints. Both systems were simulated in GROMACS 2023, with five replicates of 500 ns each.

### 2.2 Quantum chemical calculations

To evaluate the intrinsic stability of CBZ dimers independently of the protein environment, electronic structure calculations for the carbamazepine monomer and its four dimer configurations were performed using the Density Functional Theory (DFT) method in the ORCA^16^ program (version 5.0.3). Snapshots of the individual dimer configurations used in the calculations are shown in Figure S1 of the Supplementary Information. The B3LYP functional^17,18^ was selected, supplemented with the empirical D3BJ dispersion correction to properly account for non-covalent interactions. The def2-SVP basis set was used for all atoms^19^, with the RIJCOSX approximation^19–21^ applied for computational efficiency. Molecular geometries were optimized in the gas phase (vacuum) using the OPT NumFreq keywords. Solvation effects were modeled for water using the SMD (Solvation Model based on Density)^22^ continuum method.

### 2.3 Binding free-energy calculations

Relative binding free energies were estimated using the MM/GBSA approach as implemented in gmx_MMPBSA v1.6.3 ^23,24^. Calculations were performed using the Generalized Born (GB) implicit solvent model with the igb = 5 parameterization. An ionic strength of 0.15 M was applied, and the temperature was set to 310 K. Analyses were carried out on frames extracted from the joint five runs trajectory (1-5 for 2CBZ and GBM, 2-5 for 1CBZ), yielding a total of 2500 (2000) frames per system. Amber topologies were generated using the ff99SB force field for the protein and GAFF for the ligands. The MM/GBSA score was calculated as the sum of the molecular mechanics energy and solvation free energy terms:

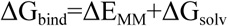

No entropic corrections were included in the MM/GBSA estimates. Reported values correspond to mean ΔG_bind_ ± standard error calculated over all analyzed frames.

Per-residue free energy decomposition was performed using the same MM/GBSA setup with idecomp = 2. The decomposition output included residues within 5 Å of the ligand (print_res = “within 5”), capturing interacting residues from both the receptor and the ligand. Per-residue contributions were decomposed into van der Waals, electrostatic, polar solvation, and nonpolar solvation terms.

### 2.4 Ligand orientation analysis

The orientational freedom of ligands within the SUR1 binding pocket was quantified by monitoring the orientation of an internal ligand vector relative to the membrane normal (Z-axis of the simulation box). For each ligand, an internal orientation vector was defined using three non-collinear atoms distributed across the molecule, selected to represent the overall ligand geometry (N1 C15 C14 for CBZ and S C21 C6 for GBM). The vector normal **n** was calculated at each simulation frame as the normalized cross product of two vectors formed by the selected atoms:

The orientation of the ligand was then characterized by the cosine of the angle θ between **n** and the Z-axis of the simulation box cos(θ)=n_z_ where n_z_ is the Z-component of the normalized vector **n**.

This definition preserves the directionality of the ligand orientation. Orientation distributions were obtained by pooling data from five independent 500 ns molecular dynamics trajectories for each ligand.

To quantify the degree of orientational ordering, the second-rank orientational order parameter S was calculated from the distribution of cos(θ):

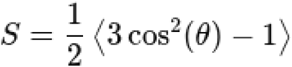

where ⟨ ⟩ denotes averaging over all frames and all trajectories.

In this definition S=0 corresponds to an isotropic orientational distribution, S>0 indicates preferential alignment along the Z-axis, S<0 indicates preferential alignment perpendicular to the Z-axis or with a defined opposite orientation.

### 2.5 Ligand density reconstruction

To enable direct comparison with cryo-EM observations, three-dimensional ligand density maps were reconstructed from MD trajectories. Prior to analysis, all frames were aligned to the SUR1 binding pocket using selected Cα atoms surrounding the ligand-binding site. The positions of all ligand atoms were then accumulated over the trajectory to generate a spatial occupancy map. Occupancy values were converted into volumetric densities and low-occupancy regions were removed using a density threshold chosen to eliminate sparsely sampled configurations. The resulting density maps represent the ensemble of ligand positions sampled during the simulations and were visualized together with the experimental cryo-EM densities using ChimeraX.

### 2.6 Trajectory analysis and visualization

We used the MDAnalysis^25^ Python library to write custom analysis scripts. For the close-contact frequency analysis, we applied a cut-off distance of 3.5 Å between atoms. Molecular visualizations were generated with ChimeraX^26^. All data and MD input files required to reproduce the results reported in this paper are available upon request.

## 3. Results

### 3.1. Intrinsic stability of CBZ dimers

The unusually large cryo-EM density observed for carbamazepine (CBZ) in the SUR1 binding pocket has previously been interpreted as potentially arising from the simultaneous binding of two CBZ molecules. To assess whether such a dimeric arrangement is intrinsically favorable, we first examined the stability of CBZ dimers using density functional theory (DFT) calculations.

Four distinct dimer configurations were considered (Figure S1), including arrangements previously reported in solution-phase studies^10^ as well as a geometry resembling the residual cryo-EM density proposed by Martin et al.^7^. Calculations were performed both in vacuum and in an implicit water environment. Relative free energies and dimerization free energies are summarized in Table S1.

The calculations indicate that CBZ molecules can form dimers that are only weakly stabilized relative to the monomeric state. The most favorable configuration corresponded to a parallel arrangement of the aromatic rings (dimer A), with a calculated dimerization free energy of approximately −2 kJ mol⁻¹. The configuration most closely resembling the shape of the unresolved cryo-EM density (dimer D) was only marginally less favorable, exhibiting a dimerization free energy close to thermodynamic neutrality (approximately +0.5 kJ mol⁻¹). The remaining configurations were substantially less stable, with positive dimerization free energies ranging from approximately 19 to 38 kJ mol⁻¹.

These results demonstrate that CBZ dimerization is chemically feasible and may occur under favorable conditions. However, the calculated stabilization energies are relatively small and do not indicate the presence of a strongly preferred dimeric state. Consequently, if a stable CBZ dimer were to exist within the SUR1 cavity, its stabilization would likely arise primarily from interactions with the surrounding protein environment rather than from intrinsic ligand–ligand interactions. To evaluate this possibility, we next investigated the behavior of both monomeric and dimeric CBZ within the SUR1 binding pocket using molecular dynamics simulations.

### 3.2. Ligand dynamics and cryo-EM density reconstruction support a dynamic single-molecule binding mode

To determine whether the cryo-EM density observed for carbamazepine is more consistent with binding of a single molecule or a CBZ dimer, we analyzed ligand dynamics within the SUR1 binding pocket using molecular dynamics simulations. As a reference, we additionally simulated glibenclamide (GBM), whose binding mode has been resolved experimentally at high resolution.

The overall stability of all simulated systems is summarized in the Supplementary Information (Figures S2–S4). Despite the removal of several intrinsically disordered regions from the model, both SUR1 and Kir6.2 remained structurally stable throughout the simulations. Having confirmed the stability of the protein scaffold, we next focused on ligand behavior within the binding pocket.

To characterize ligand mobility, we monitored the positions of representative ligand atoms over the course of the simulations (Figure 2A). For carbamazepine (CBZ), we followed the nitrogen atom within the aromatic ring, while for glibenclamide (GBM), a larger and more elongated molecule, two representative carbon atoms located at opposite ends of the ligand were monitored (marked in magenta in Figure 1E). To ensure that the observed positional variability reflected ligand dynamics rather than global protein motion, all trajectories were aligned to the SUR1 binding pocket prior to analysis. For GBM, the tracked atoms remained confined to a relatively small volume within the canonical SUR1 binding pocket. Although local fluctuations were observed, the ligand preserved a well-defined position and orientation throughout the simulations, consistent with a stable and tightly anchored binding mode.

**Figure 1.**
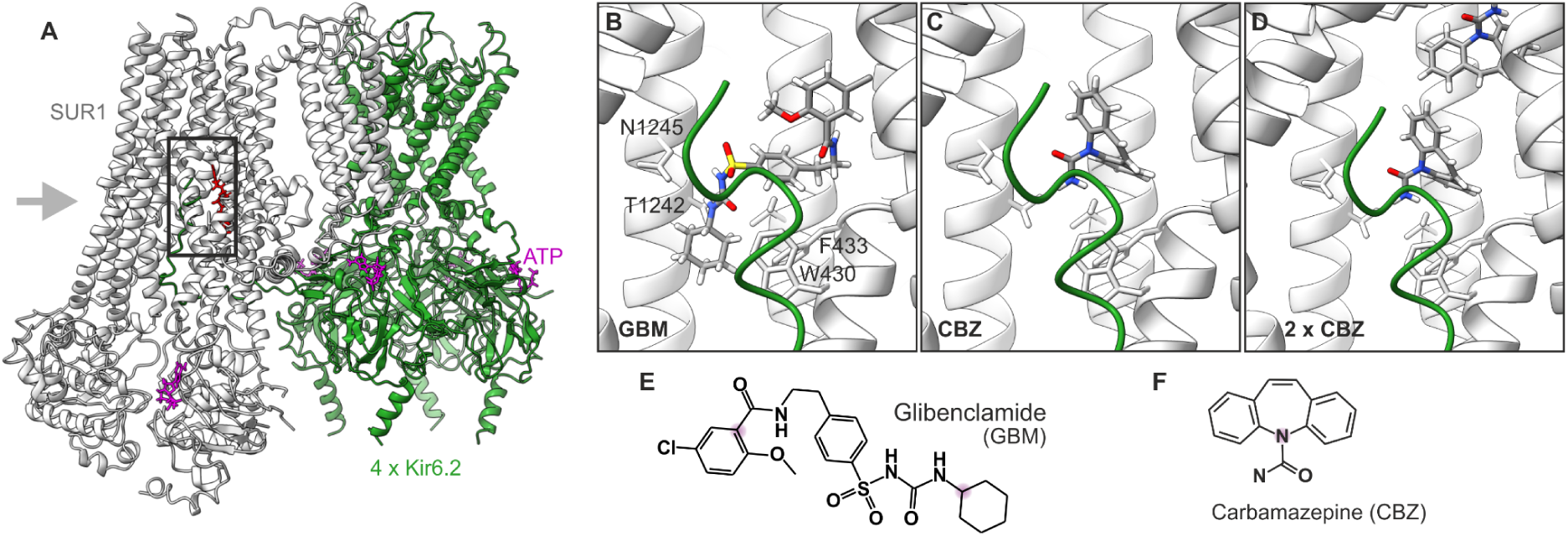
Protein system and ligand configurations used in this study. (A) The K_ATP_ channel model composed of four Kir6.2 subunits and one SUR1 subunit. ATP molecules are shown in magenta, and the high-affinity drug-binding pocket is indicated with a black box. The insets with ligands position are viewed from the grey arrow side. (B) Initial placement of glibenclamide (GBM) within the SUR1 pocket. (C) Initial configuration of a single CBZ molecule. (D) Initial configuration of the CBZ dimer, shown together with selected binding-pocket residues. (E) Chemical structure of glibenclamide. (F) Chemical structure of carbamazepine.

**Figure 2.**
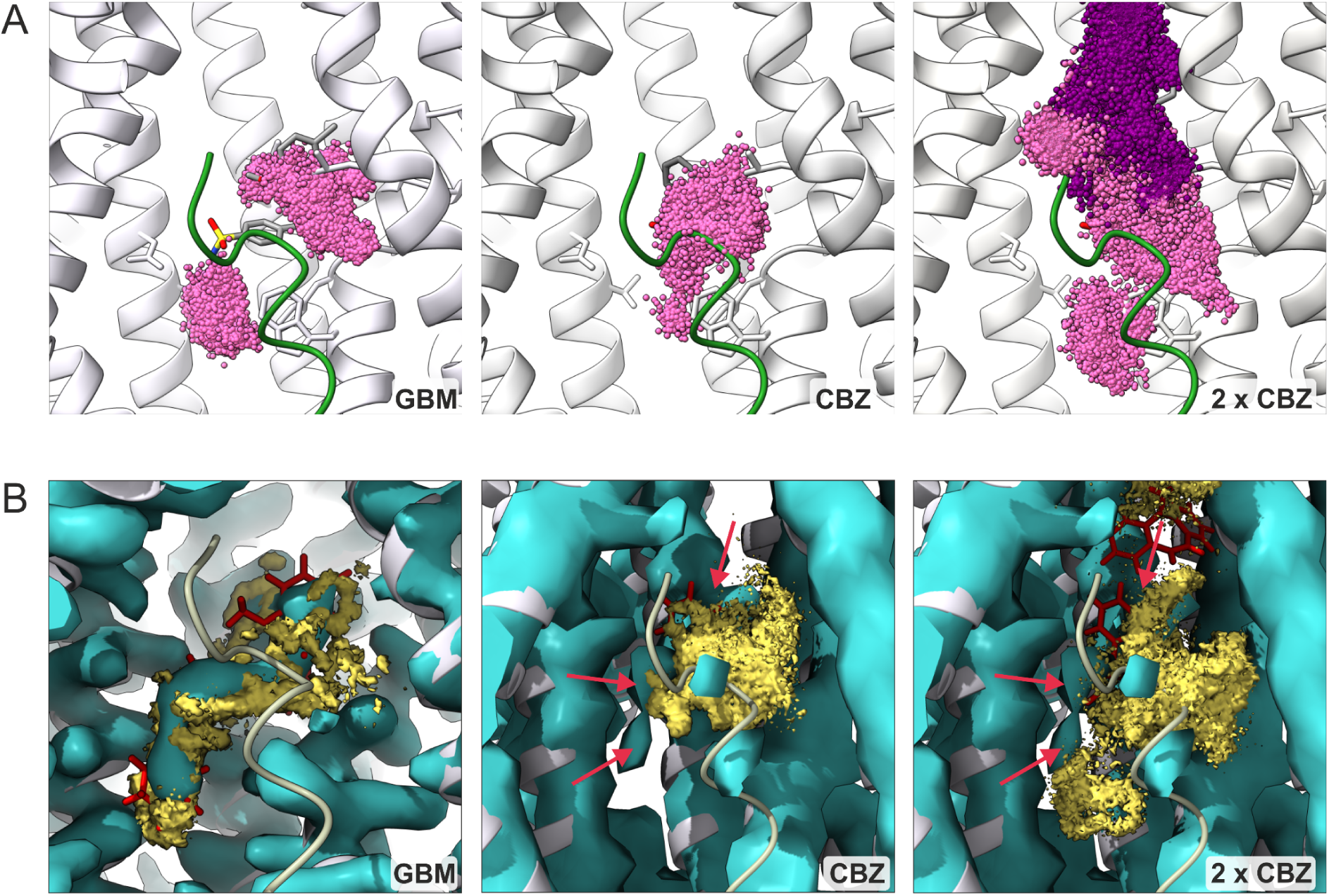
Ligand dynamics and reconstructed cryo-EM density. Top panels (A) show the positions of selected atoms sampled during simulations: two highlighted atoms for GBM (magenta), one atom for CBZ monomer (magenta), and one atom from each CBZ molecule in the dimer (magenta and violet). Bottom panels (B) display the numerically reconstructed density maps from the simulations (yellow) overlaid on the experimental cryo-EM densities (cyan). Due to the relatively low resolution of the CBZ-bound structure, additional experimental density observed by Martin et al. is indicated with red arrows for reference.

In contrast, a single CBZ molecule sampled a substantially broader region of the pocket. The ligand remained associated with the canonical drug-binding site but displayed pronounced translational and rotational motion. In four of the five independent trajectories, CBZ remained localized within the binding pocket throughout the simulation. In one trajectory, however, the ligand transiently explored deeper regions of the SUR1 cavity. Because this behavior was not reproduced in the remaining replicas, the trajectory was excluded from subsequent quantitative analyses. The corresponding distributions including this trajectory are provided in the Supplementary Information (Figure S6).

A markedly different behavior was observed when two CBZ molecules were placed simultaneously within the cavity. In this case, both ligands sampled a considerably larger volume, collectively occupying much of the accessible cavity space. Rather than forming a stable dimeric arrangement, the molecules frequently exchanged positions and competed for favorable interaction sites. This behavior suggests that the SUR1 cavity does not provide a strongly stabilizing environment for simultaneous binding of two CBZ molecules.

To directly compare the simulated ligand ensembles with experimental observations, we reconstructed three-dimensional density maps from the MD trajectories and overlaid them with the corresponding cryo-EM densities (Figure 2B). For GBM, the reconstructed density (yellow) closely reproduced the experimental cryo-EM map (cyan) in both shape and spatial extent. This agreement provides an internal validation of the simulation protocol and density reconstruction procedure.

For CBZ, the comparison proved particularly informative. The density reconstructed from simulations containing a single CBZ molecule closely matched the ligand-associated density observed in the cryo-EM structure. Despite the substantial mobility of the ligand, the sampled volume remained largely confined to the experimentally observed region. While the CBZ-bound cryo-EM structure was determined at considerably lower resolution than the corresponding GBM structure, precluding detailed atomistic comparison, the reconstructed density reproduced both the overall extent and the diffuse character of the experimental ligand-associated density. These observations indicate that the broad cryo-EM density can emerge naturally from the dynamic behavior of a single CBZ molecule within the SUR1 binding pocket.

In contrast, the density reconstructed from the two-CBZ simulations extended substantially beyond the experimental cryo-EM volume. The simulated density occupied regions of the SUR1 cavity where no corresponding experimental density is observed and partially overlapped with space sampled by the Kir6.2 N-terminal tail. Thus, while a CBZ dimer can be generated as an initial configuration, its behavior during the simulations is inconsistent with the experimentally observed density.

Taken together, these results indicate that the diffuse cryo-EM density associated with carbamazepine is more readily explained by a single dynamically bound molecule sampling multiple configurations than by the simultaneous binding of a stable CBZ dimer. The agreement between the reconstructed and experimental density maps provides direct support for a dynamic single-molecule binding mode within the SUR1 pocket.

### 3.3. CBZ binding is stabilized by a limited interaction network

The density reconstruction analysis indicates that CBZ remains localized within the SUR1 drug-binding pocket despite exhibiting substantially greater positional and orientational freedom than GBM. To identify the interactions responsible for maintaining this localization, we next analyzed residue-level contact frequencies and energetic contributions to ligand binding.

For GBM, close-contact analysis revealed a dense and highly persistent interaction network involving several SUR1 residues, most notably PHE433, ILE381, LEU434, and TRP430, all of which maintained contacts with the ligand throughout nearly the entire simulation (Figure 3). Additional interactions with TYR377 as well as residues of the Kir6.2 N-terminal tail, including ARG4 and LYS5, were also frequently observed. Together, these interactions define a well-anchored binding mode consistent with the limited mobility of GBM and the compact density observed in both simulations and cryo-EM structures.

**Figure 3:**
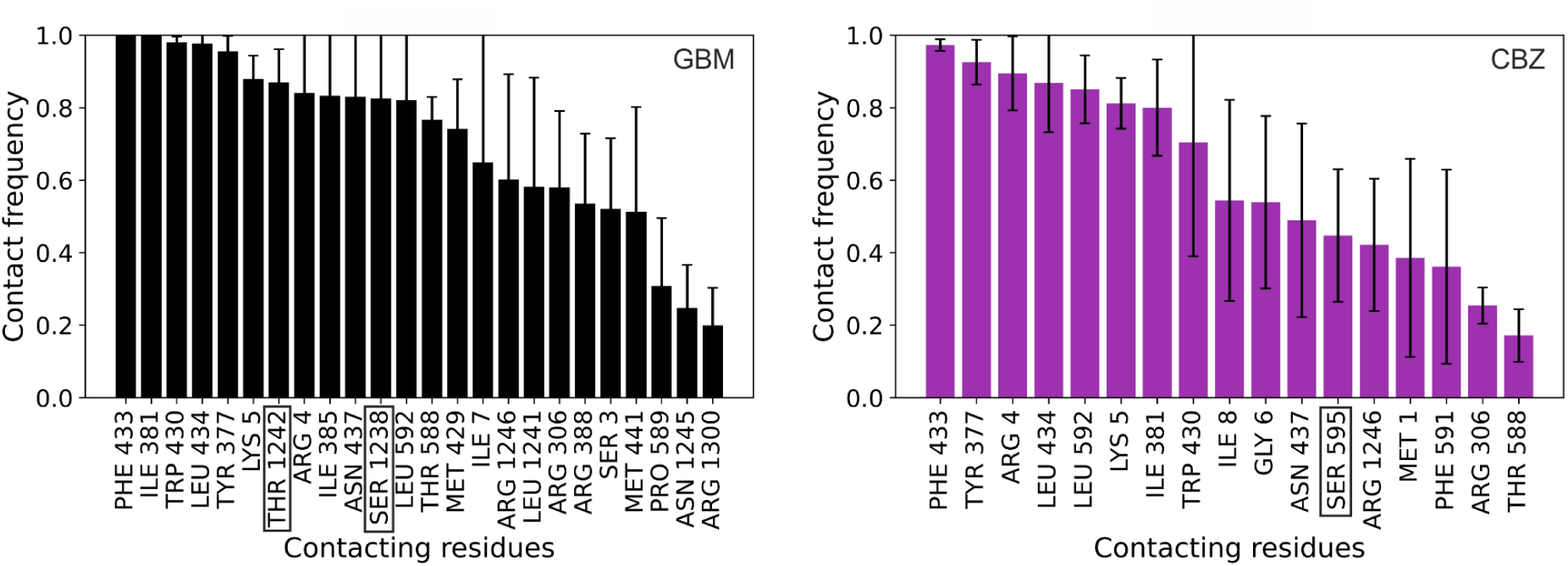
Close-contact frequencies between ligands and residues forming the SUR1 binding pocket, calculated over the MD trajectories using a 3.5 Å distance cutoff. Results are shown for glibenclamide (GBM) and carbamazepine (CBZ) in the monomeric binding mode

A markedly different interaction pattern was observed for CBZ. Although the ligand occupied the same general region of the SUR1 cavity, it formed fewer persistent contacts overall. The most prominent interaction involved PHE433, which remained in contact with the ligand for the majority of the simulation. Additional high-frequency contacts were observed with TYR377, LEU434, and ARG4, while interactions with LEU595 and LYS5 occurred more transiently. Compared with GBM, the CBZ interaction network was therefore less extensive and relied on a smaller number of frequently sampled contacts.

This interpretation is further supported by MM/GBSA per-residue energy decomposition (Figure 4). In the GBM-bound system, stabilization is distributed among several SUR1 residues, with TYR377, PHE433, and ILE381 providing the largest energetic contributions. In contrast, only a small subset of residues contributes substantially to CBZ stabilization. The strongest stabilizing contribution originates from ARG4 of the Kir6.2 N-terminal tail, while PHE433 remains the dominant contributor among SUR1 residues. Most other residues provide only minor energetic contributions, indicating that CBZ binding is maintained by a limited number of key interactions.

**Figure 4:**
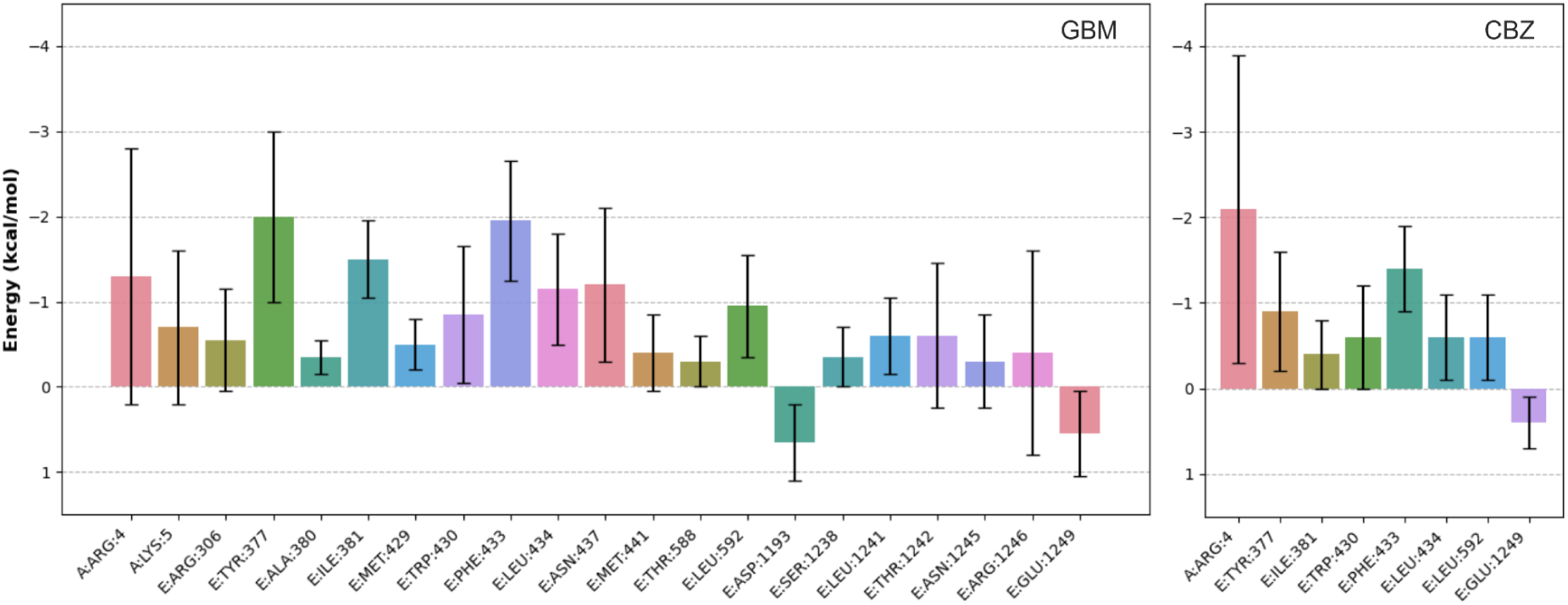
Per-residue contributions to the binding free energy of glibenclamide (GBM) and carbamazepine (CBZ), obtained from MM/GBSA calculations.

The agreement between the contact frequency and MM/GBSA analyses suggests that CBZ binding is maintained by a relatively small set of key interactions rather than by the extensive interaction network observed for GBM. In particular, recurrent interactions with PHE433 and residues of the Kir6.2 N-terminal tail appear sufficient to retain the ligand within the binding cavity while still allowing substantial translational and rotational freedom. Such a binding mode provides a straightforward molecular explanation for the broad density observed in both the MD-derived and experimental cryo-EM maps.

The two-CBZ simulations further support this interpretation. The CBZ molecule occupying the lower region of the pocket retained an interaction pattern similar to that observed for the monomeric system, including frequent contacts with PHE433 and TRP430 (Figure S5). In contrast, the second CBZ molecule failed to establish a comparable set of persistent contacts and sampled a much broader range of residues throughout the cavity. Consistently, per-residue MM/GBSA contributions for both ligands were generally weaker and less localized than in the monomeric system (Figure S8). These observations indicate that the presence of a second CBZ molecule does not generate additional stabilizing interactions and instead introduces competition for binding space within the SUR1 cavity.

### 3.4 CBZ promotes closer association with the Kir6.2 N-terminus

Both the contact frequency analysis and MM/GBSA decomposition identified recurrent interactions between CBZ and residues of the Kir6.2 N-terminal tail (KNtp), particularly ARG4 and LYS5. These observations suggest that KNtp forms an integral part of the CBZ binding environment and motivated a more detailed analysis of the spatial relationship between the ligand and the N-terminus within the SUR1 cavity.

To investigate this relationship, we reconstructed three-dimensional density maps describing the spatial distributions of both the ligand and KNtp throughout the MD trajectories (Figure 5). For GBM, the ligand density remained compact and localized within the canonical SUR1 binding pocket, consistent with its well-defined binding mode. In contrast, the density associated with KNtp was comparatively diffuse, indicating that the N-terminal tail sampled a relatively broad region of the cavity in the vicinity of the bound ligand.

**Figure 5:**
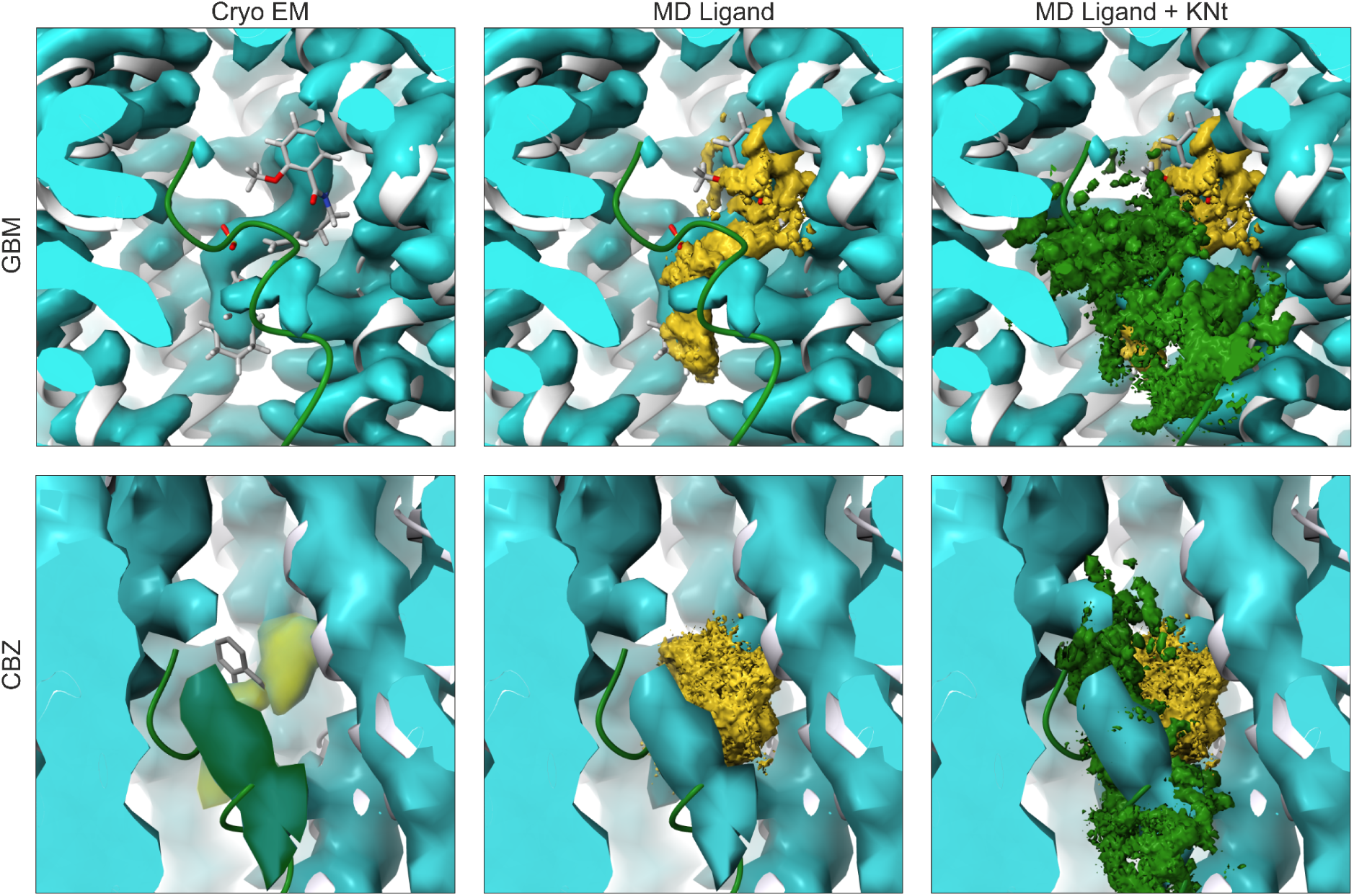
Three-dimensional density maps reconstructed from MD trajectories showing the spatial distributions of heavy atoms of the ligand and the Kir6.2 N-terminal tail (KNtp) within the SUR1 binding cavity. Density maps are shown for glibenclamide (GBM) and carbamazepine (CBZ) and are overlaid on the corresponding experimental cryo-EM density maps.

A different behavior was observed in the CBZ-bound system. Although CBZ displayed substantially greater positional variability than GBM, the ligand density remained confined to a limited region of the SUR1 cavity. At the same time, the density associated with KNtp became noticeably more localized and concentrated around the ligand. Compared with the GBM-bound system, the N-terminal tail sampled a smaller volume and remained in closer spatial proximity to the ligand throughout the simulations.

These observations are consistent with the residue-level analyses presented above, which identified frequent contacts and favorable energetic contributions involving KNtp residues. Together, the results suggest that CBZ binding is accompanied by a tighter spatial coupling between the ligand and the Kir6.2 N-terminus than observed for GBM. Given the proposed role of KNtp in stabilizing the inhibited state of KATP channels, this behavior may contribute to the mechanism by which CBZ modulates channel function and pharmacochaperone activity.

### 3.5. Orientational freedom distinguishes CBZ from GBM

The analyses presented above indicate that CBZ remains localized within the SUR1 binding pocket despite exhibiting greater positional variability than GBM. To further characterize the dynamic behavior of both ligands, we quantified their orientational freedom throughout the MD trajectories.

Ligand orientation was characterized by the distribution of cos(θ), where θ represents the angle between an internal ligand reference vector and the membrane normal (Figure 6). This analysis revealed substantial differences between the two ligands. CBZ exhibited a broad distribution spanning nearly the entire range from −1 to 1, indicating that the ligand samples a wide variety of orientations within the binding pocket. Consistent with this behavior, the orientational order parameter was close to zero (S = 0.089), characteristic of an approximately isotropic orientational distribution.

**Figure 6.**
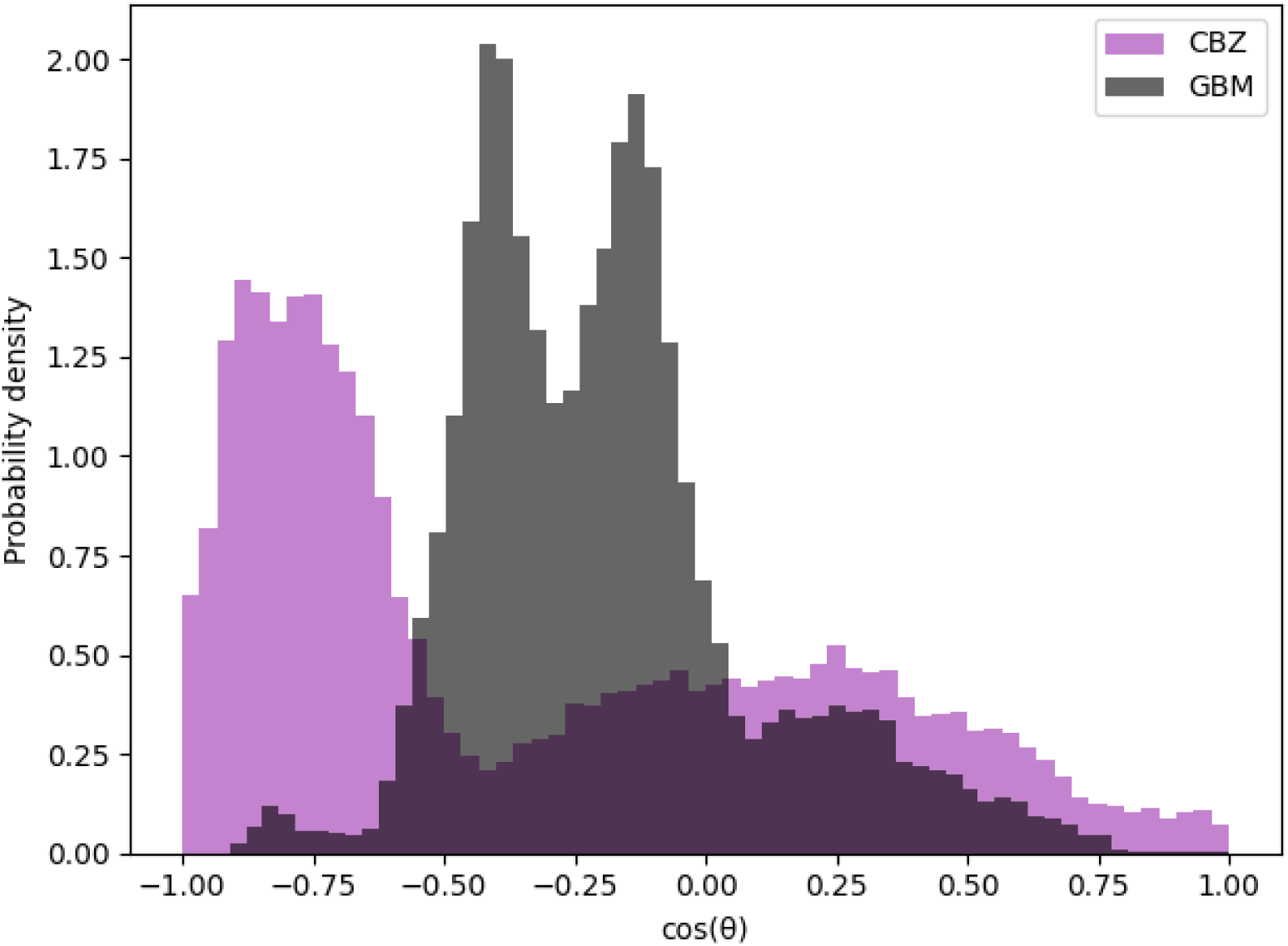
Probability density distributions of cos(𝜃) describing ligand orientation within the SUR1 binding pocket for carbamazepine (CBZ, purple) and glibenclamide (GBM, black).

In contrast, GBM displayed a considerably narrower orientational distribution, with a pronounced preference for a limited range of orientations corresponding to cos(θ) values between approximately 0 and −0.5. This orientational preference is reflected in a substantially more negative order parameter (S = −0.330), indicating a well-defined and directionally constrained binding mode.

These results complement the positional, contact-frequency, and energetic analyses presented above. While GBM is stabilized by an extensive interaction network that restricts both translational and rotational motion, CBZ remains associated with the SUR1 cavity through a smaller number of key interactions while retaining substantial orientational freedom. The ability of a single CBZ molecule to sample multiple orientations within the binding pocket provides a natural explanation for the diffuse cryo-EM density observed experimentally and is fully consistent with the density reconstruction results presented in Figure 2.

Taken together, the results support a model in which carbamazepine binds as a single, dynamically mobile ligand within the SUR1 drug-binding pocket. Rather than adopting a unique, well-defined binding pose, CBZ samples an ensemble of closely related configurations whose combined occupancy gives rise to the experimentally observed cryo-EM density.

## 4. Discussion

The interpretation of ligand-associated cryo-EM density becomes particularly challenging when ligands exhibit substantial conformational or positional heterogeneity. In such cases, the experimentally observed density represents an ensemble average over multiple molecular states rather than a single binding configuration. The CBZ-bound KATP channel structure reported by Martin et al. represents a notable example of this problem. Despite its relatively small size, CBZ produced a ligand-associated density comparable in extent to that observed for considerably larger inhibitors, leading to the proposal that either two CBZ molecules occupy the binding pocket simultaneously or that a single ligand samples multiple binding poses.

The results presented here consistently support the latter interpretation. DFT calculations indicate that CBZ dimers are only weakly stabilized in solution, suggesting that dimerization alone is unlikely to provide a strong driving force for formation of a stable dimeric complex within the confined environment of the SUR1 cavity. More importantly, molecular dynamics simulations combined with density reconstruction demonstrate that the experimentally observed cryo-EM density can be reproduced by a single CBZ molecule exhibiting substantial positional and orientational flexibility. In contrast, simulations initiated from dimeric configurations generate densities that extend beyond the experimentally observed volume and fail to reproduce the spatial characteristics of the cryo-EM map.

A key aspect of this behavior is that CBZ combines high mobility with persistent localization within the canonical SUR1 drug-binding pocket. Unlike glibenclamide, whose binding is stabilized by an extensive interaction network involving multiple SUR1 residues, CBZ relies on a more limited set of stabilizing interactions. In particular, PHE433 and residues of the Kir6.2 N-terminal tail emerge consistently across contact-frequency and MM/GBSA analyses as important contributors to ligand stabilization. This reduced interaction network allows the ligand to retain considerable translational and rotational freedom while remaining associated with the binding cavity.

The involvement of the Kir6.2 N-terminal tail is particularly noteworthy. Previous structural studies have suggested that KNtp contributes to ligand-mediated channel inhibition by occupying the SUR1 cavity adjacent to the drug-binding site. Our simulations extend this picture by showing that CBZ is associated with a tighter spatial coupling to KNtp than observed for GBM. Although the present simulations do not directly address channel gating or pharmacochaperone activity, the observed interactions between CBZ and KNtp suggest a possible structural basis for the functional role of the N-terminus in CBZ-mediated channel regulation.

More broadly, these findings highlight the importance of considering molecular ensembles when interpreting ligand-associated cryo-EM densities. For small and conformationally flexible ligands, experimentally observed densities may arise not from a single well-defined binding mode but from the superposition of multiple closely related configurations. In such cases, integrating molecular dynamics simulations with density reconstruction provides a useful framework for distinguishing between alternative structural interpretations and relating experimental densities to underlying molecular behavior.

## 5. Conclusions

In summary, our results demonstrate that the diffuse cryo-EM density associated with carbamazepine in the SUR1 drug-binding pocket is best explained by a single dynamically bound ligand rather than by the formation of a stable CBZ dimer. While CBZ exhibits substantially greater positional and orientational freedom than glibenclamide, it remains localized within the binding cavity through a limited set of stabilizing interactions involving both SUR1 residues and the Kir6.2 N-terminal tail. The close agreement between experimentally observed and MD-derived density maps highlights the importance of ligand dynamics in the interpretation of cryo-EM data and provides a molecular explanation for the unusual density observed in CBZ-bound KATP channel structures.

## Acknowledgements

The authors thank Show-Ling Shyng and Camden Driggers for valuable discussions and insights regarding KATP channel structure and pharmacological regulation. KWS acknowledged Polish high-performance computing infrastructure PLGrid for awarding this project access to the LUMI supercomputer, owned by the EuroHPC Joint Undertaking, hosted by CSC (Finland) and the LUMI consortium through PLL/2025/09/018870. We also thank Interdisciplinary Centre for Mathematical and Computational Modelling ICM, University of Warsaw, for awarding computational resources under Grant GA76-10. Christina Karafyllia’s participation in this project was supported by the Toruń Students Summer Program (TSSP), Nicolaus Copernicus University in Toruń, Poland.

## Supplementary Information

### 1. DFT Calculations of Carbamazepine Dimer Stability

**Figure S1:**
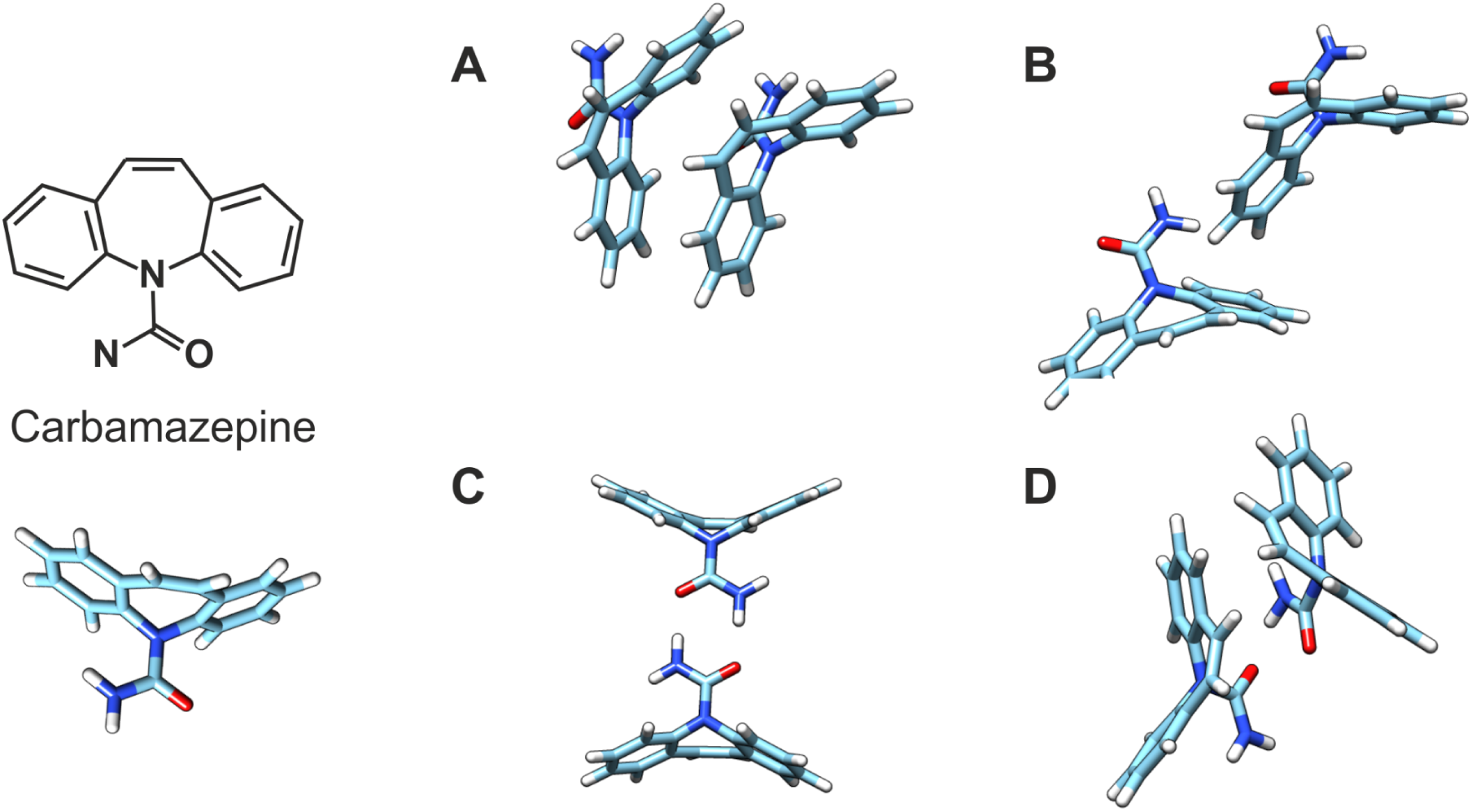
Initial carbamazepine (CBZ) structures used in the DFT calculations. The upper panel shows the optimized geometry of a single CBZ molecule. Panels A–D present the four dimer configurations examined in this study after geometry optimization in water using ORCA. Configuration C corresponds to the dimer arrangement previously reported in crystal structures and solution-phase studies of CBZ dimers. Configuration D was designed to resemble the arrangement proposed by Martin et al. as a possible explanation for the unresolved ligand-associated cryo-EM density observed in the SUR1 binding pocket. Relative energies and dimerization free energies for all configurations are reported in Table S1.

**Table S1:**
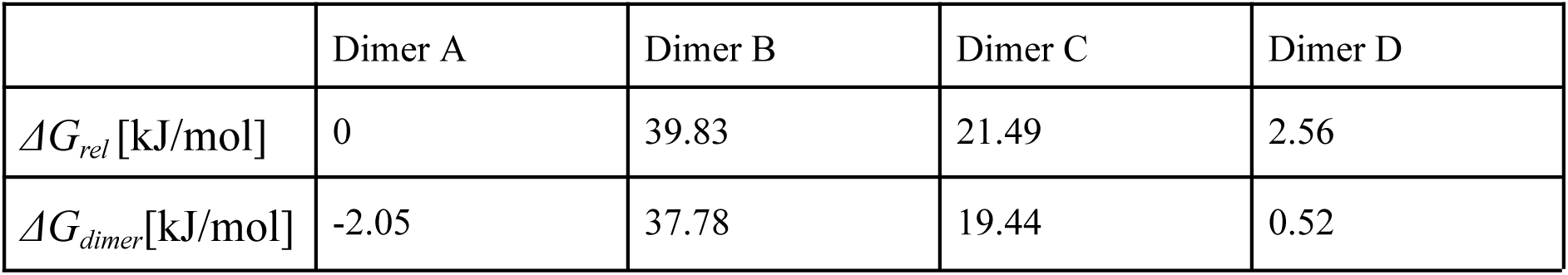
ORCA-calculated relative Gibbs free energy of dimer stability (ΔG_rel_), referenced to the most stable configuration, and the Gibbs free energy of dimerization (ΔG_dimer_), calculated as the difference between the dimer’s free energy and the doubled free energy of the monomer.

### 2. Molecular Dynamics simulations stability

The stability of the protein system throughout the simulations was assessed by calculating the Root Mean Square Deviation (RMSD), determined separately for the Kir6.2 and SUR1 subunits (Figure S1). Prior to analysis, all trajectories were aligned to the vicinity of the ligand binding site within the upper parts of the TMD1 and TMD2 domains. Specifically, alignment was performed using the Cα atoms of residues CYS6, PHE305, LEU366, LEU434, VAL587, VAL1250, GLU1253, and ASN1301, which are located on the α-helices encompassing the ligand(s). This alignment strategy might have slightly increased the calculated RMSD values for more distant parts of the complex, such as Kir6.2 and the NBD domains of SUR1. Nevertheless, the RMSD of Kir6.2 did not exceed 4 Å, plateauing at an average of 3 Å for all systems (see Figure 1). For the SUR1 protein, the RMSD exhibited minor fluctuations, plateauing at an average of about 3.5 Å. For 2 CBZ, the SUR1 RMSD values were slightly higher, and they occasionally peaked above 4 Å. To identify the protein regions responsible for these elevated RMSD values, we performed an additional Root Mean Square Fluctuation (RMSF) analysis for SUR1. This analysis revealed that the higher RMSD values observed in the SUR1 were primarily attributed to the dynamic movements of the NBD domains, particularly NBD2, as well as the terminal ends of the unmolded IDR (Intrinsically Disordered Region) segments.

**Figure S2:**
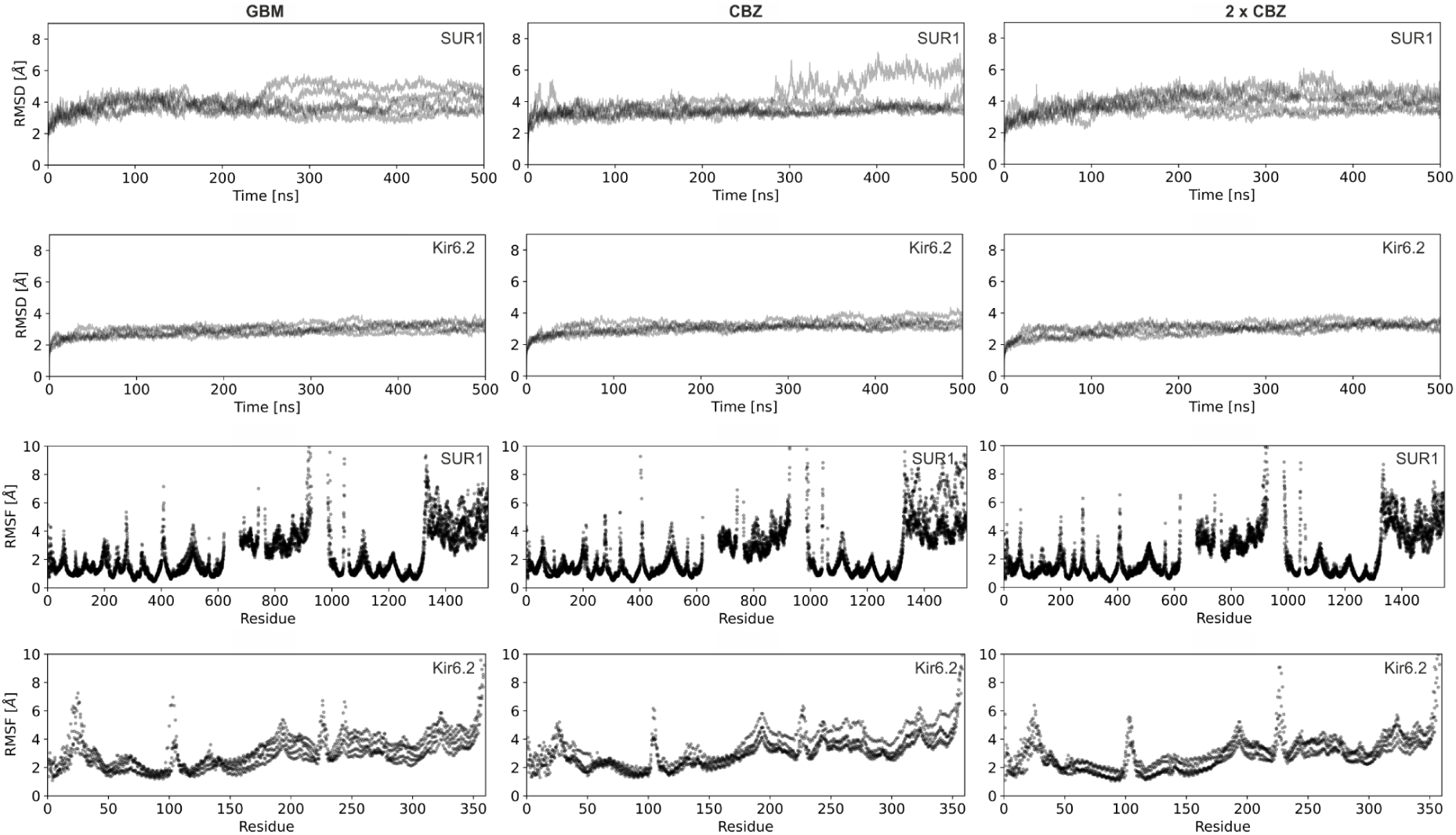
Root mean square deviation for each system shown for Kir6.2 tetramer and SUR1. Root mean square fluctuations for SUR1 (left to right: 1 CBZ, 2 CBZ, GBM).

### 3. 4xKir6.2-1xSUR1 system dynamics

To further characterize the behavior of the protein complex during the simulations, we analyzed several geometric parameters of the system (figure 2). We calculated the “propeller angle,” which describes the orientation of SUR1 relative to the Kir6.2 subunits, and the distance between the NBD domains. The propeller angle in most available structures of pancreatic KATP channels ranges from 45 to 55 degrees, indicating that our systems are consistent with the known conformational variability of the pancreatic KATP channel structure. The inter-domain distance varies from 35 to 43 Å (fully inward-open SUR1 conformations), and its probability peaks combine to form a broad one.

**Figure S3:**
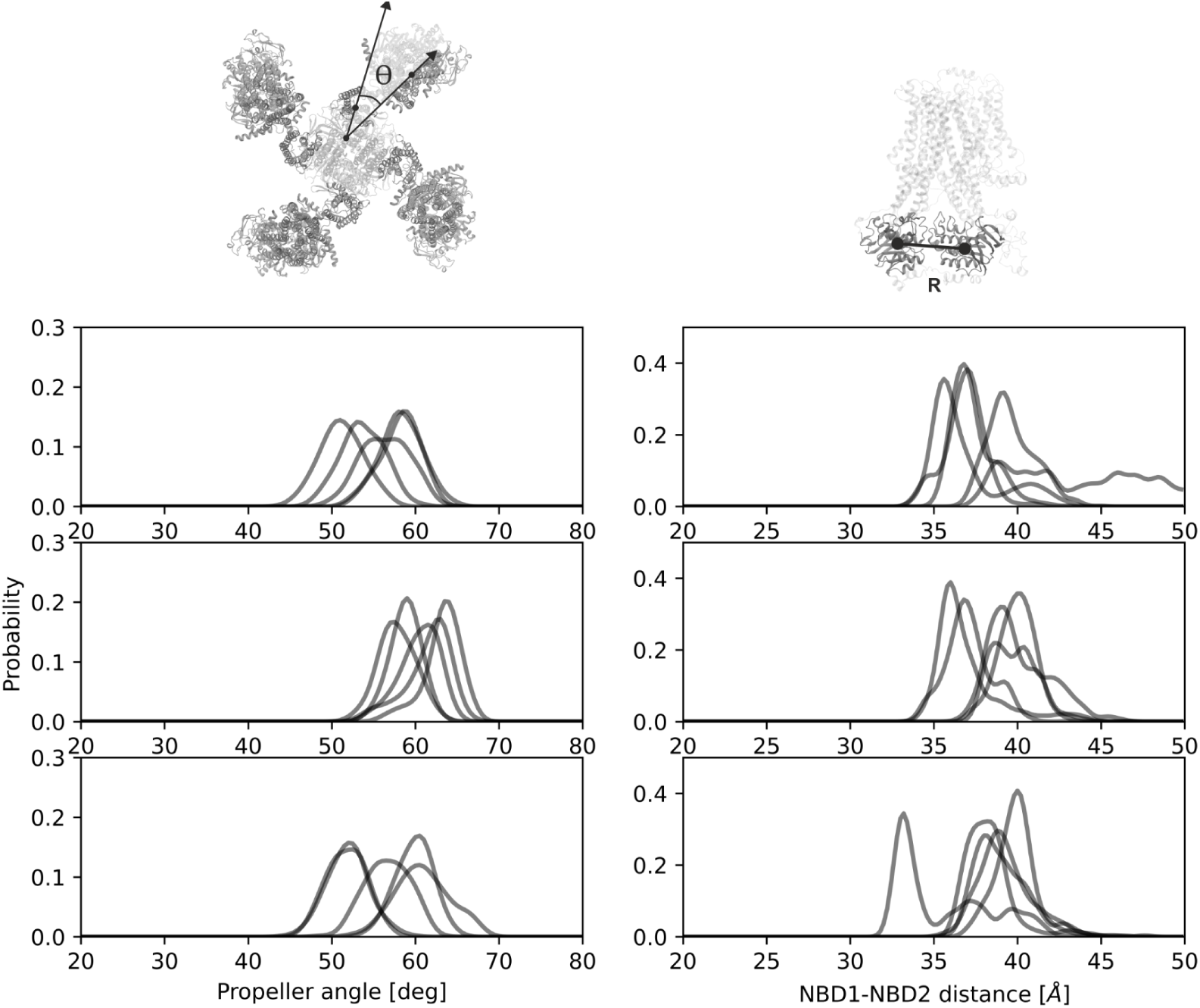
Distributions of the propeller angle and the distance between nucleotide-binding domains (NBD1 - NBD2) calculated over the MD trajectories. The top panels show results for the system containing one carbamazepine molecule (1×CBZ), the middle panels for the system containing two carbamazepine molecules (2×CBZ), and the bottom panels for the glibenclamide-bound system (GBM). Data are shown separately for individual simulation replicas, illustrating the range of conformations sampled during the simulations.

### 4. Ligand Binding within the SUR1 Pocket

**Figure S4:**
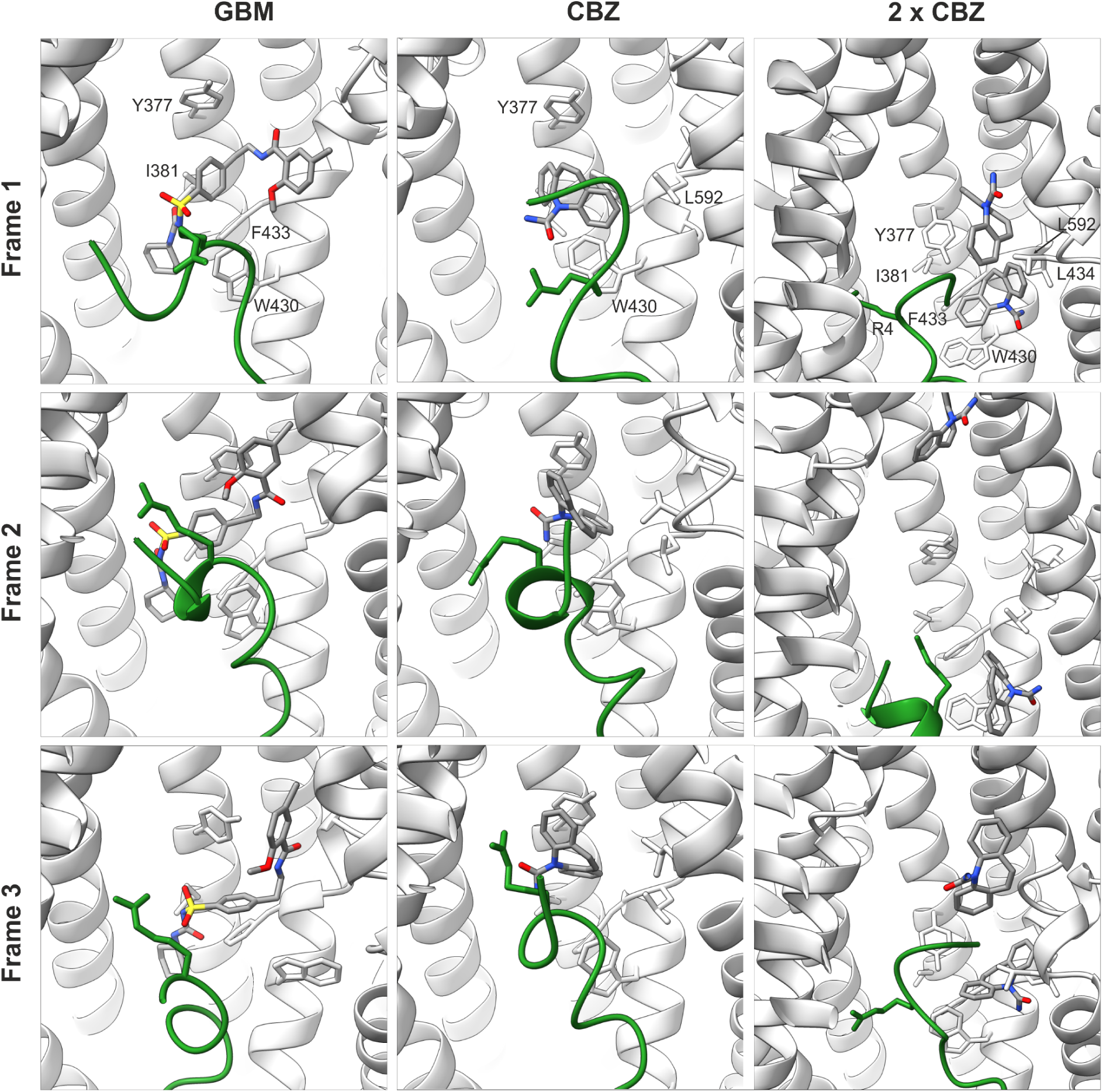
Representative snapshots extracted from molecular dynamics simulations illustrating ligand binding within the SUR1 pocket. The left column shows three snapshots from independent trajectories of the glibenclamide-bound system (GBM), the middle column shows snapshots from the system containing one carbamazepine molecule (1×CBZ), and the right column shows snapshots from the system containing two carbamazepine molecules (2×CBZ). The Kir6.2 N-terminal tail (KNtp) is shown in green in all panels.

**Figure S5:**
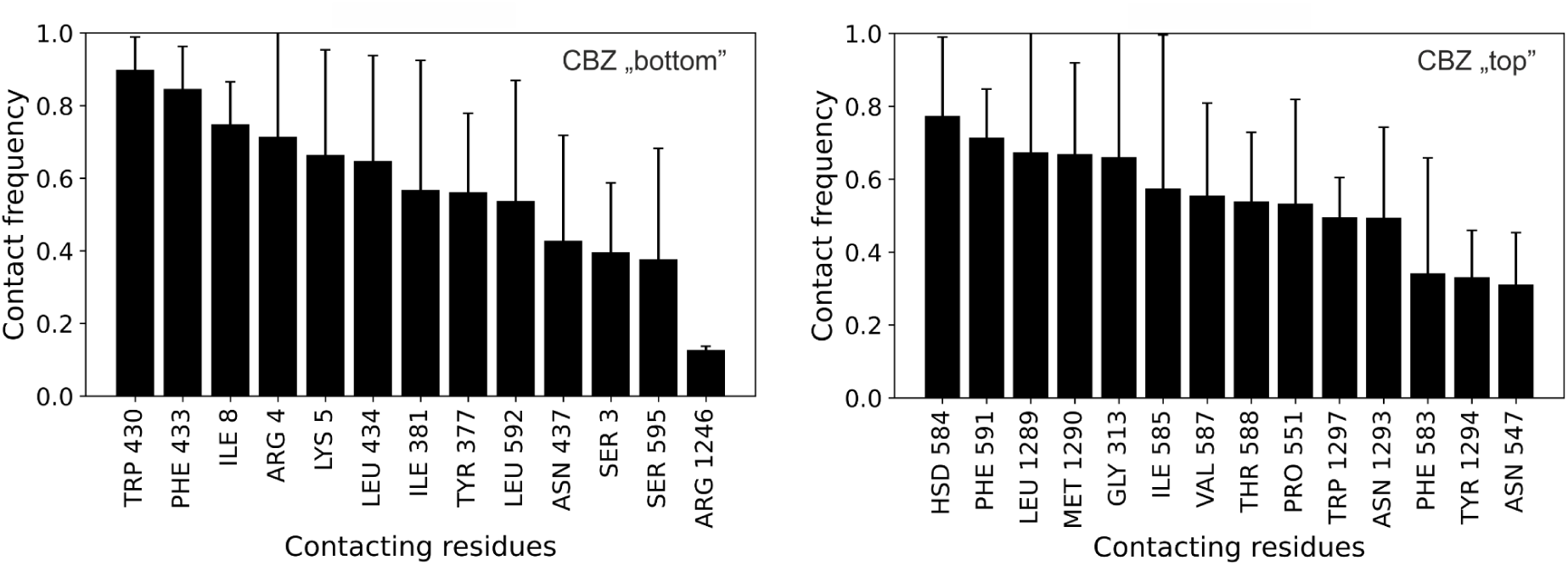
Close-contact frequencies between carbamazepine molecules and residues forming the SUR1 binding pocket for simulations containing two CBZ molecules. Results are shown separately for the bottom and top CBZ molecule. Contacts were defined using a 3.5 Å distance cutoff.

**Table S2:**
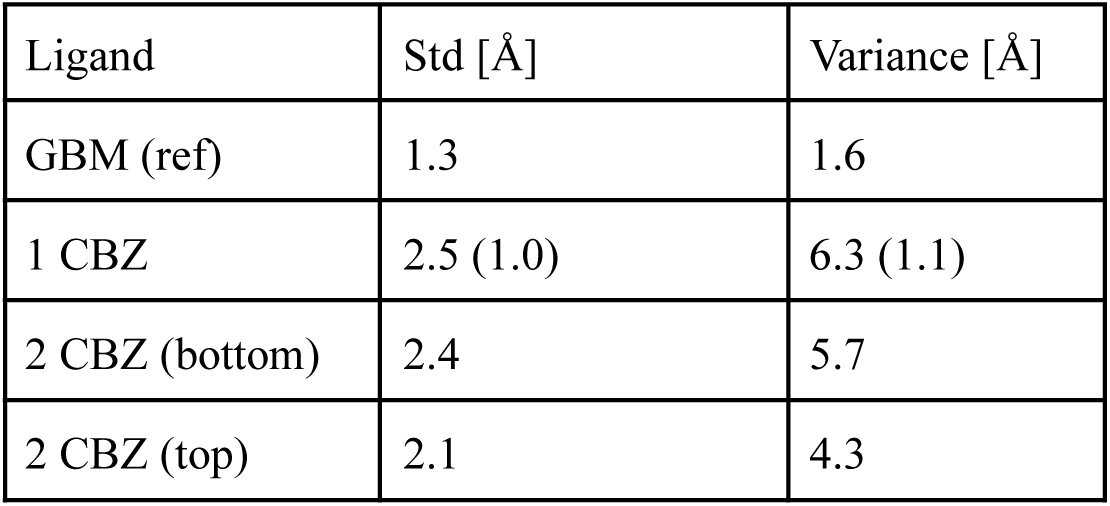
Standard deviation (STD) and variance of ligand atomic positions calculated relative to their mean positions over the MD trajectories. For the 1×CBZ system, values reported in parentheses correspond to calculations performed after excluding trajectory 1, in which the ligand transiently exits the canonical binding site.

**Figure S6:**
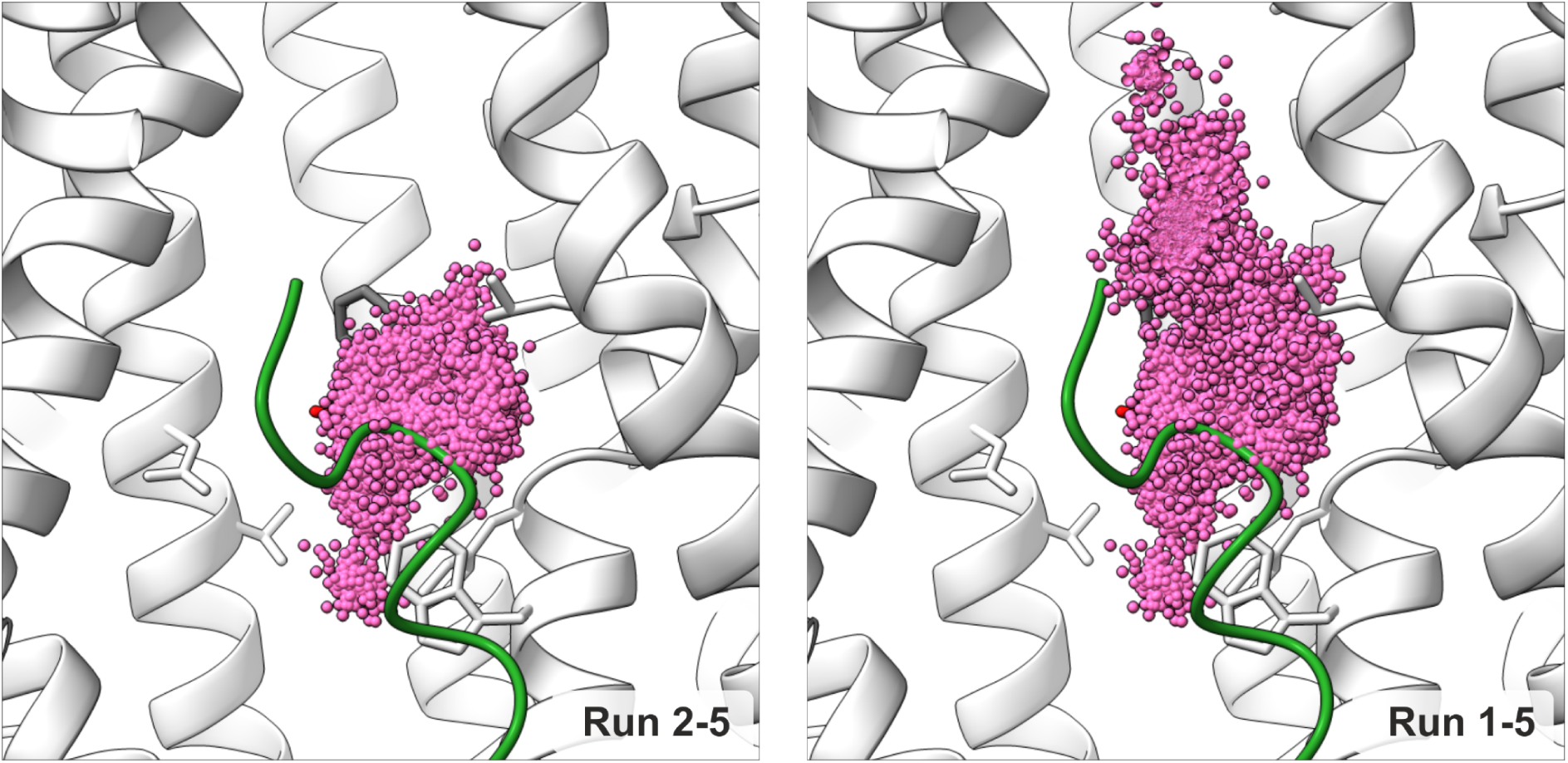
Distributions of CBZ atomic positions projected onto the protein structure. The left panel shows positional distributions calculated excluding trajectory 1, in which CBZ transiently exits the canonical binding site, while the right panel shows distributions including this trajectory. The comparison illustrates the impact of rare exploratory events on the overall positional spread.

**Figure S7:**
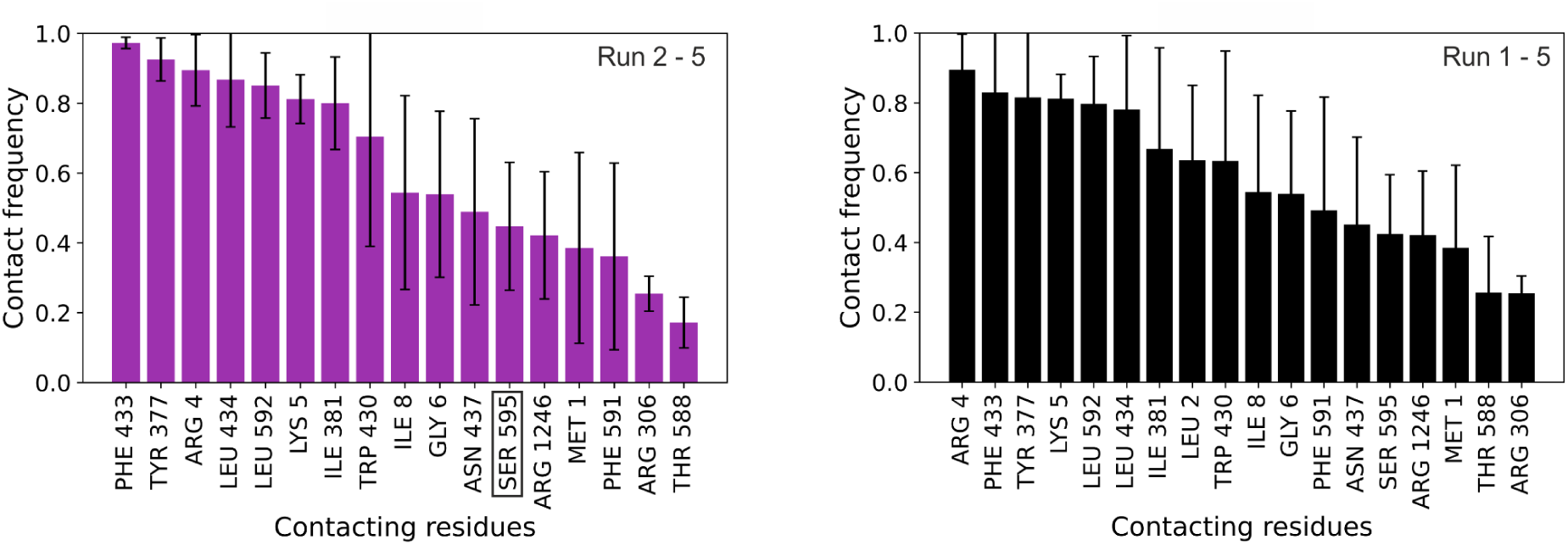
Close-contact frequency distributions for the CBZ monomer system calculated excluding (left) and including (right) trajectory 1, in which the ligand transiently explores deeper regions of the SUR1 cavity. This comparison highlights the influence of ligand escape events on contact statistics.

**Figure S8:**
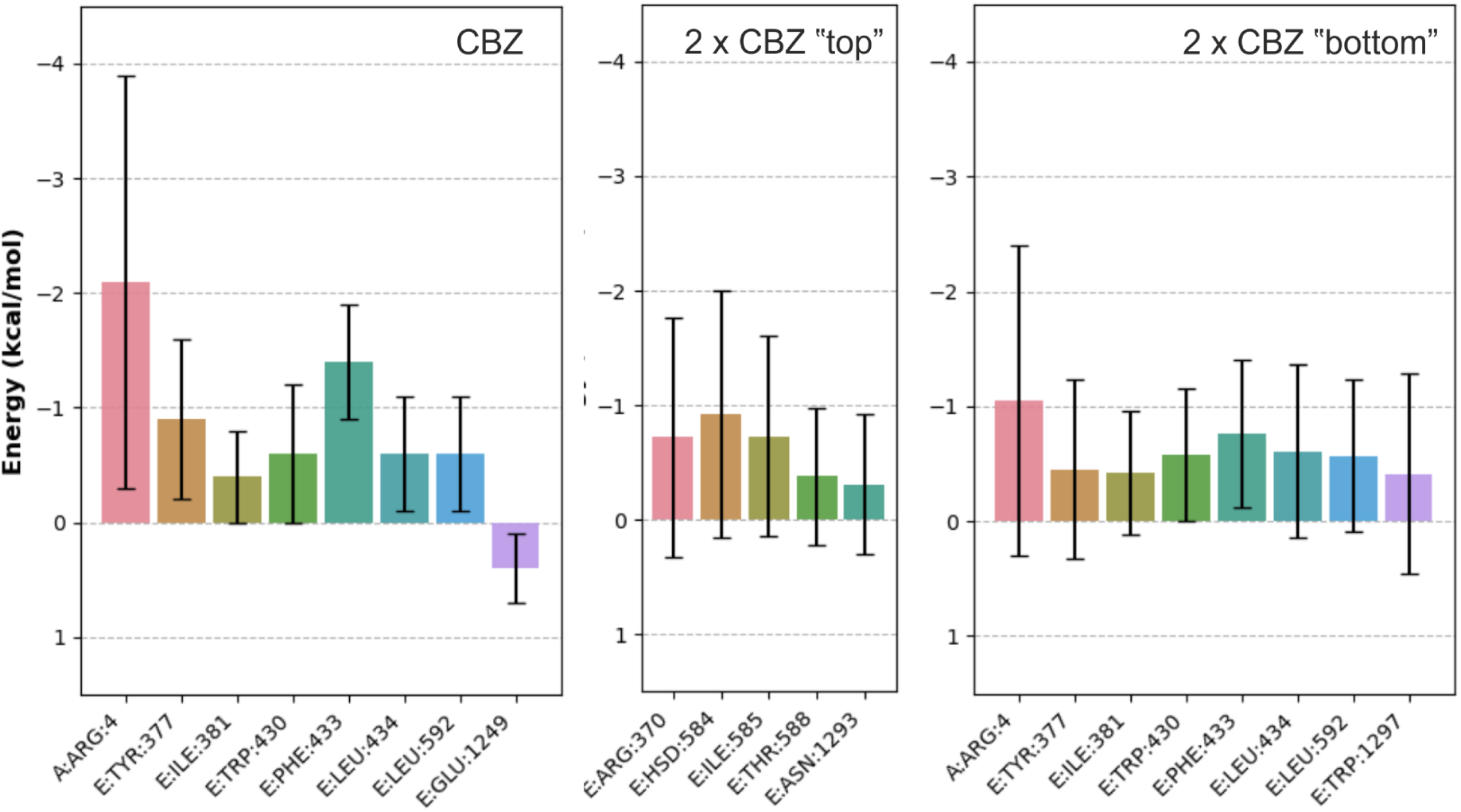
Per-residue MM/GBSA binding free energy contributions for the CBZ monomer system compared with those obtained for the two-CBZ simulations. Results are shown separately for the CBZ molecule located deeper in the pocket (“top”) and the second CBZ molecule (“bottom”), illustrating differences in residue-level energetic contributions between the binding modes.

## Notes

### Competing Interest Statement

The authors have declared no competing interest.

## References

1. Patton, B. L., Zhu, P., ElSheikh, A., Driggers, C. M. & Shyng, S.-L. Dynamic duo: Kir6 and SUR in K channel structure and function. Channels (Austin) 18, 2327708 (2024).

2. Puljung, M. C. Cryo-electron microscopy structures and progress toward a dynamic understanding of K channels. J Gen Physiol 150, 653–669 (2018).

3. Driggers, C. M. & Shyng, S.-L. Mechanistic insights on KATP channel regulation from cryo-EM structures. J Gen Physiol 155, (2023).

4. ElSheikh, A. et al. Identification and rescue of congenital hyperinsulinism-associated ABCC8 mutations that impair K channel trafficking. J Biol Chem 301, 110779 (2025).

5. Pipatpolkai, T., Usher, S., Stansfeld, P. J. & Ashcroft, F. M. New insights into K channel gene mutations and neonatal diabetes mellitus. Nat Rev Endocrinol 16, 378–393 (2020).

6. Martin, G. M. et al. Mechanism of pharmacochaperoning in a mammalian K channel revealed by cryo-EM. Elife 8, (2019).

7. Martin, G. M., Kandasamy, B., DiMaio, F., Yoshioka, C. & Shyng, S.-L. Anti-diabetic drug binding site in a mammalian K channel revealed by Cryo-EM. Elife 6, (2017).

8. Zhou, Q. et al. Carbamazepine inhibits ATP-sensitive potassium channel activity by disrupting channel response to MgADP. Channels (Austin) 8, 376–382 (2014).

9. Wu, J.-X. et al. Ligand binding and conformational changes of SUR1 subunit in pancreatic ATP-sensitive potassium channels. Protein Cell 9, 553–567 (2018).

10. Hunter, C. A., McCabe, J. F. & Spitaleri, A. Solvent effects of the structures of prenucleation aggregates of carbamazepine. CrystEngComm 14, 7115–7117 (2012).

11. Hall, A. V., Cruz-Cabeza, A. J. & Steed, J. W. What Has Carbamazepine Taught Crystal Engineers? Crystal Growth & Design (2024) doi:10.1021/acs.cgd.4c00555.

12. Sung, M. W. et al. Ligand-mediated Structural Dynamics of a Mammalian Pancreatic K Channel. J Mol Biol 434, 167789 (2022).

13. PubChem. Carbamazepine. https://pubchem.ncbi.nlm.nih.gov/compound/2554.

14. Wu, E. L. et al. CHARMM-GUI Membrane Builder toward realistic biological membrane simulations. J Comput Chem 35, 1997–2004 (2014).

15. Abraham, M. J. et al. GROMACS: High performance molecular simulations through multi-level parallelism from laptops to supercomputers. SoftwareX 1-2, 19–25 (2015).

16. Neese, F., Wennmohs, F., Becker, U. & Riplinger, C. The ORCA quantum chemistry program package. J Chem Phys 152, 224108 (2020).

17. Stephens, P. J., Devlin, F. J., Chabalowski, C. F. & Frisch, M. J. Ab initio calculation of vibrational absorption and circular dichroism spectra using density functional force fields. J. Phys. Chem. 98, 11623–11627 (1994).

18. Becke, A. D. Density-functional thermochemistry. III. The role of exact exchange. J. Chem. Phys. 98, 5648–5652 (1993).

19. Weigend, F. & Ahlrichs, R. Balanced basis sets of split valence, triple zeta valence and quadruple zeta valence quality for H to Rn: Design and assessment of accuracy. Phys Chem Chem Phys 7, 3297–3305 (2005).

20. Weigend, F. Accurate Coulomb-fitting basis sets for H to Rn. Phys Chem Chem Phys 8, 1057–1065 (2006).

21. Neese, F., Wennmohs, F., Hansen, A. & Becker, U. Efficient, approximate and parallel Hartree–Fock and hybrid DFT calculations. A ‘chain-of-spheres’ algorithm for the Hartree–Fock exchange. Chem. Phys. 356, 98–109 (2009).

22. Marenich, A. V., Cramer, C. J. & Truhlar, D. G. Universal solvation model based on solute electron density and on a continuum model of the solvent defined by the bulk dielectric constant and atomic surface tensions. J Phys Chem B 113, 6378–6396 (2009).

23. Valdés-Tresanco, M. S., Valdés-Tresanco, M. E., Valiente, P. A. & Moreno, E. gmx_MMPBSA: A New Tool to Perform End-State Free Energy Calculations with GROMACS. Journal of Chemical Theory and Computation (2021) doi:10.1021/acs.jctc.1c00645.

24. Miller, B. R., 3rd et al. MMPBSA.py: An Efficient Program for End-State Free Energy Calculations. J Chem Theory Comput 8, 3314–3321 (2012).

25. Michaud-Agrawal, N., Denning, E. J., Woolf, T. B. & Beckstein, O. MDAnalysis: a toolkit for the analysis of molecular dynamics simulations. J Comput Chem 32, 2319–2327 (2011).

26. Meng, E. C. et al. UCSF ChimeraX: Tools for structure building and analysis. Protein Sci 32, e4792 (2023).

